# Functional Metagenomic Analysis Reveals Early Gut Microbiota Alterations in Alpha-Synuclein Transgenic Mice: Insights into Parkinson’s Disease Progression

**DOI:** 10.1101/2025.03.18.644066

**Authors:** Divya Mondhe, Nanthini Jayabalan, Man Tamang, Jessika Suescun, Richard Gordon

**Affiliations:** School of biomedical science, Faculty of health, Queensland University of Technology, Brisbane, Australia; Translational Research Institute, Brisbane, Australia; UQ Centre for Clinical Research, University of Queensland, Australia

## Abstract

Parkinson’s Disease (PD) is the fastest growing neurodegenerative disease and manifests as a synucleinopathy with progressive motor and non-motor symptoms. In this study, we investigated the gut-microbial alterations associated with PD in an α-synuclein transgenic mouse model (hα-Syn) at both early and late stages of the disease. We utilised high-resolution functional metagenomics with ultra-deep sequencing to ensure the identification of low-abundance and novel taxa. The microbial community and metabolic pathways were profiled using Microba community and pathway profiler, respectively. While the microbial alpha-diversity remained unchanged between the hα-Syn and WT group across different disease stages, distinct shifts in microbial composition were observed. The hα-Syn group form two separate clusters corresponding to early and late stages of the disease, indicating progressive dysbiosis. Gut dysbiosis in the early stages of PD was characterised with an increase in *Staphylococcus* species and a decrease of *Duncaniella* and *Muribaculum* species. A reduction in lactobacillus genera was also observed in PD. Furthermore, microbes associated with SCFA production declined whereas and opportunistic pathogens increased in abundance. These findings provide evidence supporting the hypothesis that microbiota alterations may contribute to the onset and progression of PD, highlighting potential microbial targets for future therapeutic interventions.

## Introduction

Parkinson’s disease (PD) is the second most common, chronic, progressive, neurodegenerative disease worldwide. Non-motor symptoms (NMS) in Parkinson’s disease (PD) often emerge decades before the onset of motor symptoms, significantly impacting patients’ quality of life. These early symptoms include gastrointestinal dysfunction, sleep disturbances, mood disorders, cognitive impairment, and autonomic dysfunction [7, 8]. Among them, constipation, is consider particularly a common prodromal sign, sometimes appearing 10–20 years before motor symptoms become evident. The presence of these early NMS suggests that PD pathology may begin outside the central nervous system, particularly in the gut and olfactory pathways, before spreading to the substantia nigra [10, 11]. Alterations in the gut microbiome composition (dysbiosis) associated with PD have been linked to significant changes in metabolic functions and disease progression. Evidence from α-synuclein mouse models suggests that the gut microbiome may influence the spread of α-synuclein from the gut to the brain via the vagus nerve [13]. This supports the idea that PD pathology may originate in the gut and later spread to the CNS. Studies involving vagotomy in α-synuclein models have shown that severing this communication pathway reduces the spread of α-synuclein from the gut to the brain, further implicating the gut microbiota in the initiation of PD pathology[45, 46].

Studies using α-synuclein transgenic mouse models have provided compelling evidence linking gut dysbiosis to neuroinflammation, α-synuclein aggregation, and motor dysfunction, offering insights into how gut-brain axis alterations may contribute to PD progression[13, 14]. Cohorts studies have identified at least 44 gut microbial species that differ in abundance between people with PD (PwPD) and healthy individuals [15].

Notably, members of genera *Akkermansia, Bifidobacterium and Lactobacillus* are consistently increased in PD[16-20], while*Roseburia, Faecalibacterium* and *Blautia* are significantly reduced [20, 21]. Genera *Roseburia, Faecalibacterium* are particularly important due to their role in production of short chain fatty acids (SCFAs). SCFA play essential role in maintenance of intestinal homeostasis[22]. In PD, an increased abundance of opportunistic pathogens and microbes associated heightened intestinal permeability is observed, alongside a reduction in anti-inflammatory microbial species [23].

Predictive functional analysis of shotgun sequenced gut microbiome suggest that microbial metabolism and downstream metabolic pathways are dysregulated in PD. A recent Meta-analysis identified disruptions in synthesis and metabolism of neuroactive molecules, including dopamine, gamma amino butyric acid and serotonin, further implicating the gut microbiome in PD pathophysiology [24].

In this study we aimed to characterize gut microbial changes associated with Parkinson’s disease (PD) progression using the widely stidied A53T α-synuclein overexpressing M83 transgenic (hαSyn-Tg) mouse model. This model express the human A53T α-synuclein variant under the mouse prion protein promoter (PrP)[25-27] and develops severe motor impairments around 12 months of age without overt neuronal loss. Early motor impairment is marked by wobbling and posturing followed by complete loss of movement at later stages. To investigate the relationship between gut microbiota composition and disease progression, we performed a longitudinal analysis of fecal microbiota in hαSyn-Tg and wild-type (WT) mice at both pre- and post-onset stages of motor symptoms. By identifying age- and α-synuclein-dependent microbial alterations, this study provides insights into potential gut-derived mechanisms contributing to PD pathogenesis.

## Materials and methods

### Animals

The A53T α-Synuclein overexpressing M83 line (hαSyn-Tg), expresses the full length A53T variant of human α-Synuclein under a mouse prion promoter[25-27]. hαSyn-Tg mice were bred by crossing homozygous and hemizygous mice. The hαSyn-Tg mice were originally developed in the laboratory of John Trojanowski, Dept. Pathology and Laboratory Med, University of Pennsylvania and were procured from the JAX laboratories (Bar harbour, Maine, US). The transgenic mice were housed at the animal facility at The University of Queensland, Brisbane, Australia. The research was approved by the Institutional Animal ethics committee (AEC) at the University of Queensland.

### Faecal Pellet collection

Faecal pellets were collected from all mice at two timepoints (9-11 weeks, pre-onset of motor symptoms and ---weeks, post onset of motor symptoms). For faecal pellets collection, individual mice were transferred into a sterile bowl and were allowed to defecate, and the fresh faecal pellets were collected in a sterile microcentrifuge tube using a sterile toothpick and samples were immediately placed on dry ice. The faecal samples were stored at −80°C until further processing for gut microbiome analysis.

### Metagenomics sequencing

DNA was extracted from the faecal sample. DNA concentration was determined using nanodrop. The quality of the extracted DNA was assessed using qubit and was used to prepare libraries for shotgun sequencing. The prepared libraries were sequenced on Illumina NovaSeq™ 6000 at Microba.

### Bioinformatics Analysis

The sequence reads were analysed using proprietary bioinformatic Microba Analysis Platform (MAP). This platform consists of the Microba Genome Database (MGDB), the Microba Genes Database (MGENES), the Microba Community Profiler (MCP), and the Microba Genes and Pathway Profiler (MGPP).

#### Taxonomic Profiling

Taxonomic profiling was performed using the MCP. MCP quantifies the relative abundance of species within a metagenomic sample using the MGDB as a genome reference. MGDB is Microba’s high-quality and expansive genome database, collated and curated from public repositories such as GenBank, and supplemented with faecal derived species that Microba has mined from public and private metagenomic data. The Genome Taxonomy Database (GTDB) is used as the primary source of taxonomic annotation of the resulting representative genomes (http://gtdb.ecogenomic.org/). Profiling is performed after all data are quality controlled. The results of the MCP are both the relative abundances of species, as well as read counts assigned to each species. Reads that cannot be identified are listed as “Unmapped” (the read had no high-quality alignment in the database), or “Unassigned” (the read had a high-quality alignment, however there was insufficient evidence to determine that the species was present). Reads may remain unassigned because the species they originated from was present at very low abundance, or because complex strain heterogeneity results in inadequate statistical confidence for the presence of a species. The MCP is parameterised conservatively and has been validated to report less than 1% false positive species.

#### Functional Profiling

Microba Gene and Pathway Profiler (MGPP) was used for functional profiling of the sequenced reads. The Microba gene functional data is also used to quantify higher level metabolic pathways. These pathway annotations are provided based on the MetaCyc database (https://metacyc.org/). The MetaCyc pathway abundance pipeline summarises an input Enzyme Commission (EC) matrix into pathways (modular metabolic pathways such as glycolysis), and metabolic groups (groups of pathways that share a role in metabolism e.g. Carbohydrate metabolism). For the quantification of pathways in the metagenomic samples, each genome is annotated with the fraction of all genes present in all pathways in MetaCyc (based on EC annotations). Pathways that are complete or near complete (completeness > 80%), are stored as a reference database for further analysis. Following this metabolic group abundances are quantified as the summed abundance of the pathways within each group.

### Statistical analysis

All statistical analyses were performed using R (version 4.3.3)[28].

The microbial abundance tables and their associated metadata were imported as data frames in R.

#### Alpha diversity indices

To determine the intra sample microbial diversity within the hαSyn-Tg and WT groups, alpha diversity indices (Species richness and Shannon diversity) were calculated using *vegan* package in R[29]. The species richness indicates the total number of microbial species present in a sample or community, whereas Shannon diversity index takes the abundance of microbial species along with their presence/absence to determine the diversity within a sample. It is calculated as, *H* = −Σ*p*_i_ * *ln(p*_i_) where *p*_i_ is the proportion of the entire community made up by species *i*.

#### Beta diversity indices

To determine the inter sample differences between the hαSyn-Tg and WT mice beta-diversity analysis was performed using the *vegan* package in R[29]. To determine the dissimilarities in the microbial composition between the hαSyn-Tg and WT mice, *bray-curtis dissimilarity matrix* were made, using the *vegdist package*. To visualise the pattern of community composition, principal coordinate analysis (PCoA) and non-metric multidimensional scaling (NMDS) were performed using the *cmdscale* and *metaMDS* functions respectively. Permutational multivariate analysis of variance (PERMANOVA) analysis was performed to determine the statistical significance of the observed differences using the *adonis2* function[30].

#### Differential abundance analysis

*DESeq2* and *Maaslin2* R packages were used to determine the differences in the gut microbial abundance between transgenic and WT mice[31, 32]. Both *DESeq2* and *Maaslin2* were used with default set of parameters. For *DESeq2* analysis non-normalised count datasets were used. *DESeq2* tests differential expression using negative binomial generalized linear models by performing normalisation using relative log expression. *Maaslin2* tests for differential expression in normalised data using gaussian linear models. The data is then normalised using the total sum scaling. Both the methods use Wald’s tests to find the statistical significance and associated p-values.

#### Visualisation

All the analysis performed were visualised using the *ggplot2* package in R[33].

## Results

### Synuclein accumulation is accompanied by early changes in the gut microbiota

Taxonomic profiling of the gut microbiome from α-synuclein A53T transgenic (hαSyn-Tg, n=8) and wild type (WT, n=6) mice at early stages (3weeks of age) identified a total of 40 microbial species across both hαSyn-Tg and WT groups. An overlap of 27 microbial species was observed across both the groups (Figure S1). Furthermore,7 and 10 microbial species were unique to hαSyn-Tg and WT groups respectively. A non-significant decrease (p>0.05) in the gut microbial alpha diversity indices (species richness and Shannon diversity index) were observed in the hαSyn-Tg (n=8) group compared to the WT (n=6) (Figure 1A and 1B). The transgenic hαSyn-Tg (n=8) and WT (n=6) occupied two independent clusters (PERMANOVA, R^2^=0.25097, F=4.0207 and p-value < 0.05) in principal coordinate analysis (Figure 1C), demonstrating differences in the gut microbiome composition between the transgenic and WT mice. The first two principal components, PCoA1 and PCoA2 explained 32% and 28% of the total variances, respectively indicating that most variation in the microbiome could be captured by these two components. The PCoA2 showed a positive correlation with the WT mice. Similarly, NMDS analysis identified two independent clusters for the hαSyn-Tg (n=8) and WT (n=6) with a stress value of 0.109 (Figure 1D). The gut microbiome at early stages of synuclein accumulation in hαSyn-Tg mice and the WT is dominated by phylum *Firmicutes A* (hαSyn-Tg 47.7%, WT 24.1%) *and Bacteroidota* (hαSyn-Tg 18.4%, WT 31.3%). The hαSyn-Tg mice showed a higher Firmicutes/ Bacteroides ratio (F/B ratio=1.08), compared to the WT mice (F/B ratio = 0.34) (Figure S2). At genera level, *genus COE1* from the *Lachnospiraceae* family dominates the gut microbiome of the hαSyn-Tg mice (n=8) and the WT (n=6). The gut microbiome in the WT (n=6) is predominantly composed of the genera *Muribaculum* (17.73%) and *Lactobacillus* (12.2%) although the hαSyn-Tg mice (n=8) show no presence of the genus *Muribaculum* and a lowered abundance of genus *Lactobacillus*. Gut microbial species *COE1 sp002358575, CAG-485 sp002362485, Lactobacillus_B animalis* form 70% of the microbiome in the hαSyn-Tg mice (n=8), whereas in the WT (n=6) COE*1 sp002358575, Muribaculum intestinale, Lactobacillus johnsonii, Dubosiella newyorkensis and CAG-485 sp002362485* form the majority of the gut microbiome. Differential abundance analysis identified *14-2 sp000403315, ASF356 sp000364165, Kineothrix sp000403275, Staphylococcus nepalensis, Staphylococcus xylosus, UBA1394 sp900066845, Hungatella A sp002358555* gut microbial species to be significantly enriched (p<0.05) the hαSyn-Tg mice (n=8) with the WT (n=6) as reference (Figure 1F). Gut microbial species such as *Bifidobacterium pseudolongum A, COE1 sp000403215, Dubosiella newyorkensis, Muribaculum intestinale, Duncaniella muris, Parasutterella excrementihominis, Muribaculum sp002473395* and *Lactobacillus johnsonii* to be signifacntly depleted (p<0.05) in the hαSyn-Tg mice (n=8) (Figure 1F).

**Figure 1:**
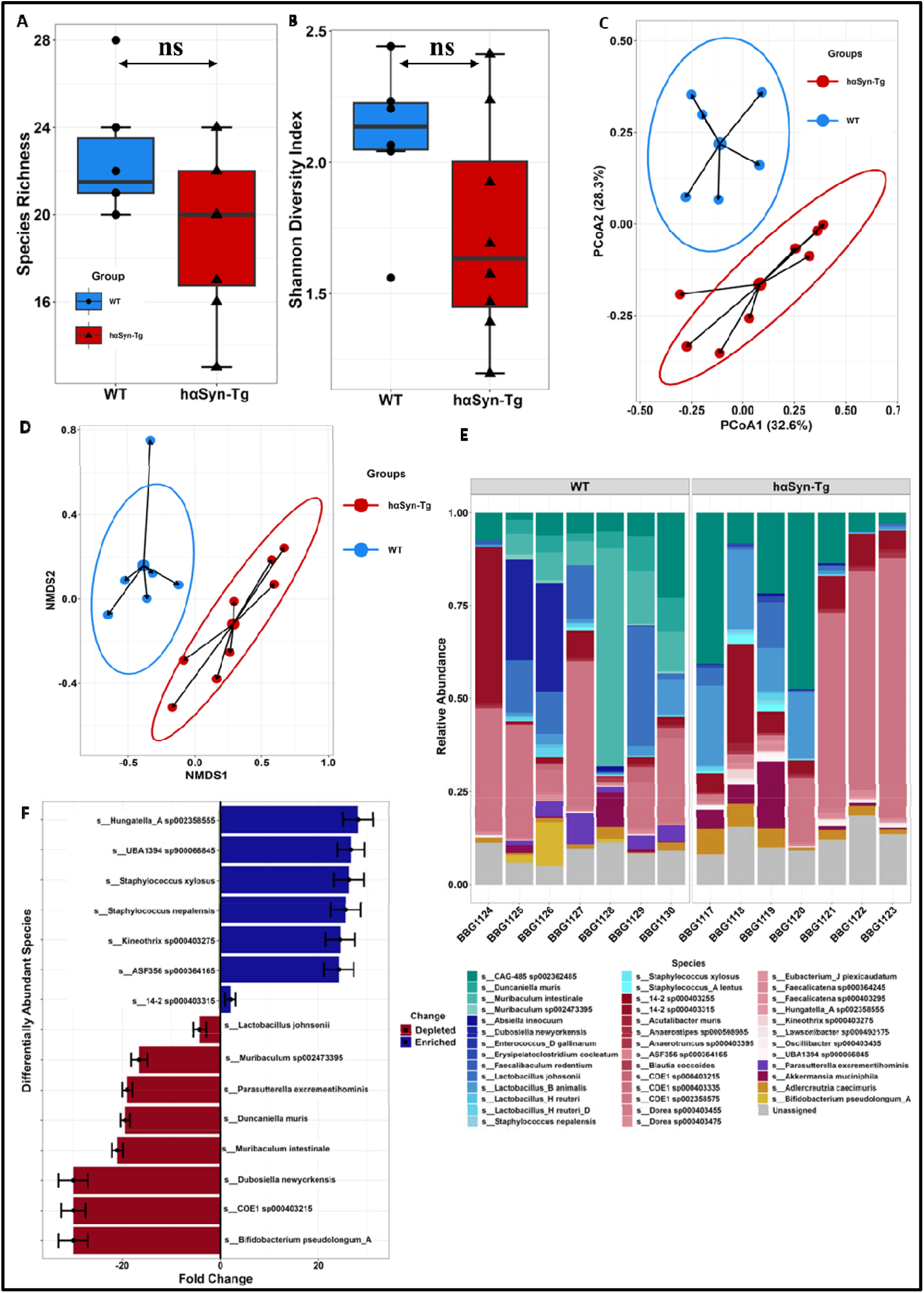
Synuclein accumulation is accompanied by early changes in the gut microbiota. Box plot highlighting the differences in the alpha diversity measures A) Species Richness and B) Shannon diversity index between the gut microbiome of Wild type (WT, n=6) and α-Synuclein transgenic (hαSyn-Tg, n=8) mice pre-onset of motor symptoms. C) Principal coordinate analysis of the gut microbiome by WT (n=6) and hαSyn-Tg (n=8) mice pre-onset of motor symptoms, using the bray-curtis dissimilarities. D) NMDS analysis of the gut microbiome by WT (n=6) and hαSyn-Tg (n=8) mice pre-onset of motor symptoms. E) Taxonomic relative abundance of gut microbial species across by WT (n=6) and hαSyn-Tg (n=8) mice pre-onset of motor symptoms. F) Differential abundance of gut-microbial species in hαSyn-Tg (n=8) compared to WT (n=6) mice using DEseq2.

### Prolonged gut dysbiosis with synuclein pathology

The hαSyn-Tg mice prior to motor deficits around 7-8 months of age start showing αSynuclein inclusions in selected neuronal populations including the mid-brain, brainstem, and cerebellum. Faecal metagenomic analysis of hαSyn-Tg mice (n=8) and WT (n=8) at 9 months of age revealed gut microbial changes post onset of motor symptoms in the hαSyn-Tg mice. A total of 50 gut microbial species were identified in the hαSyn-Tg and the WT mice post onset of motor symptoms. Both the groups showed an overlap of 37 microbial species (Figure S1). 7 gut microbial species were unique to the hαSyn-Tg (n=8) group, and 6 microbial species were unique to the WT (n=8) group. No significant differences were observed in the alpha diversity indices (species richness and Shannon diversity index) between the hαSyn-Tg and the WT mice (Figure 2A and 2B). Principal coordinate analysis revealed that WT and the hαSyn-Tg mice form overlapping clusters suggesting similar composition of the gut microbiome across groups post-onset of motor symptoms (PERMANOVA, R2=0.02861, F=0.4123 and p-value >0.05) (Figure 2C). PCoA1 accounts for 34.5% of the total variance in the dataset and shows a positive correlation with the hαSyn-Tg and WT gut microbiomes. Similarly, NMDS analysis revealed an overlap between the microbes of the hαSyn-Tg and WT, with a stress of 0.141 (Figure 2D). The gut microbiome post-onset of motor symptoms is dominated by the phylum *Bacteroidota* (WT: 59.9% and hαSyn-Tg:53.3%) and *Firmicutes A* (WT: 19.5% and hαSyn-Tg: 25.7%). A non-significant change in the F/B ratio was observed between the WT (F/B ratio = 0.117) and the hαSyn-Tg (F/B ratio = 0.177) groups. At genera level genera *Duncaniella, CAG-485 (family: Muribaculaceae)* dominate the gut microbiome of the WT and hαSyn-Tg. The genera *Akkermansia* is highly abundant in the hαSyn-Tg mice (6.6%) whereas in the WT, its abundance is limited to 0.7% post-onset of motor symptoms. Post onset of motor symptoms, in the hαSyn-Tg mice *CAG-485 sp002362485, Duncaniella muris, COE1 sp002358575* and *Akkermansia munciniphila* form approximately 70% of the total gut microbiome, whereas in the WT *CAG-485 sp002362485, Duncaniella muris, Muribaculum intestinale* and *14-2 sp002358575* form the majority of the gut microbiome (Figure 2E). Differential abundance analysis of the gut microbiome post onset of motor symptoms identified *Parasutterella excrementihomonis*, *Adlercreutzia mucosicola, Lactobacillus H reuteri, Lactobacillus johnsonii* and *14-2 sp000403255* to be significantly depleted (p>0.05) in the hαSyn-Tg (n=8) mice with the WT (n=8) as reference. Simulatneously, *Flavonifractor plautii* and *Clostridium Q symbiosum* were significantly enriched (p>0.05) in the hαSyn-Tg mice post onset of the motor symptoms (Figure 2F).

**Figure 2:**
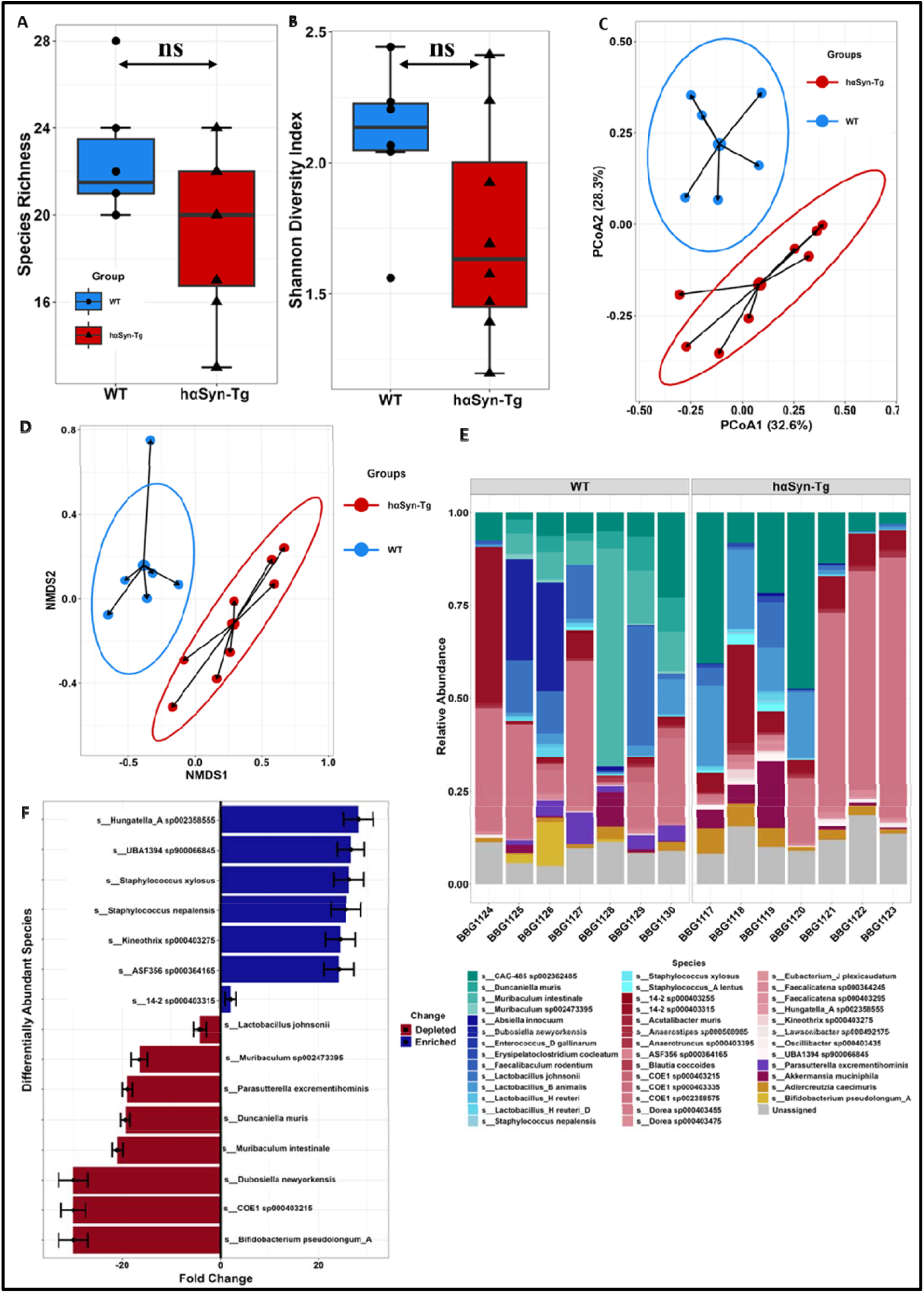
Prolonged gut dysbiosis with synuclein pathology. Box plot highlighting the differences in the alpha diversity measures A) Species Richness and B) Shannon diversity index betweenthe gut microbiome of Wild type (WT, n=8) and α-Synuclein transgenic (hαSyn-Tg, n=8) mice post-onset of motor symptoms. C) Principal coordinate analysis of the gut microbiome by WT (n=8) and hαSyn-Tg (n=8) mice pre-onset of motor symptoms, using the bray-curtis dissimilarities. D) NMDS analysis of the gut microbiome by WT (n=8) and hαSyn-Tg (n=8) mice pre-onset of motor symptoms. E) Taxonomic relative abundance of gut microbial species across by WT (n=6) and hαSyn-Tg (n=8) mice pre-onset of motor symptoms. F) Differential abundance of gut-microbial species in hαSyn-Tg (n=8) compared to WT (n=8) mice using DESeq2.

### Both ageing and synuclein pathology influence the gut microbiota

Changes in the gut microbial composition are influenced by both the presence of the transgene and age of the mice (Figure 3). Microbial species such as *Akkermansia munciniphila* and *UBA1394* are negatively associated with the presence of the transgene and show a weak positive correlation with the age, i.e., These microbes deplete in the presence of the transgene and show a non-significant increase with age. Members of the *Lactobacillus genera (Lactobacillus H reuteri, Lactobacillus johnsonii* and *Lactobacillus B animalis)* show a negative correlation with age. These are found to be depleted in the gut as the mice age, whereas in the hαSyn-Tg mice these are enriched in abundance. A similar trend is also observed in the members of the *genera Staphylococcus*. Certain microbial species such as *Muribaculum* and *Duncaneilla muris* show a positive correlation with both ageing and the presence of the transgene. These microbes are found to be enriched in the hαSyn-Tg mice with age. *Parasutterella excrementhihominis* shows a positive correlation with genotype and a negative correlation with age, as a result it is enriched in the hαSyn-Tg mice and depletes in abundance with age. *Clostridium Q symbiosum* on the other hand shows a positive correlation with the age and a negative correlation with the genotype, allowing it to be enriched in abundance with age and deplete in the presence of the transgene.

**Figure 3:**
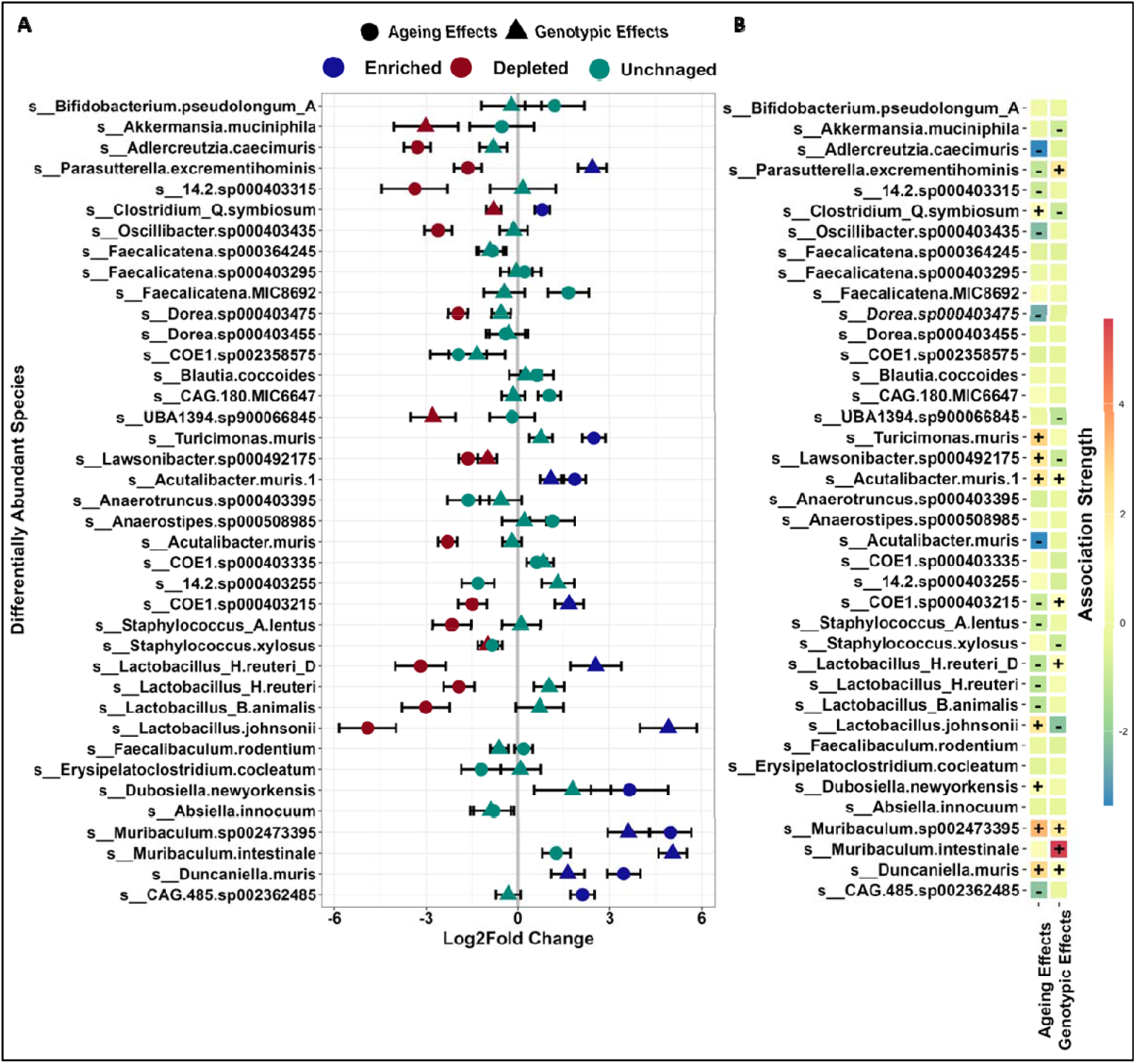
Both ageing and synuclein pathology influence the gut microbiota. A) Differential abundance plot highlighting gut microbial species being altered due to ageing and genotype in hαSyn-Tg mice. The circles and the squares represent the microbial alterations due to ageing and genotypic effects respectively. Blue and red colour represents microbes significantly enriched (p-value <0.05) and significantly depleted (p-value <= 0.05) in hαSyn-Tg mice as compared to the WT. Green color represents microbial changes that are statistically non-significant (p value>0.05)in the hαSyn-Tg mice as compared to the WT. B) Heat-map of the correlation between the gut microbial species abundance in hαSyn-Tg mice with ageing and genotypic effects. Positive and negative significant microbial correlations (p-values <0.05) have been indicated with + and – signs respectively.

**Figure 4:**
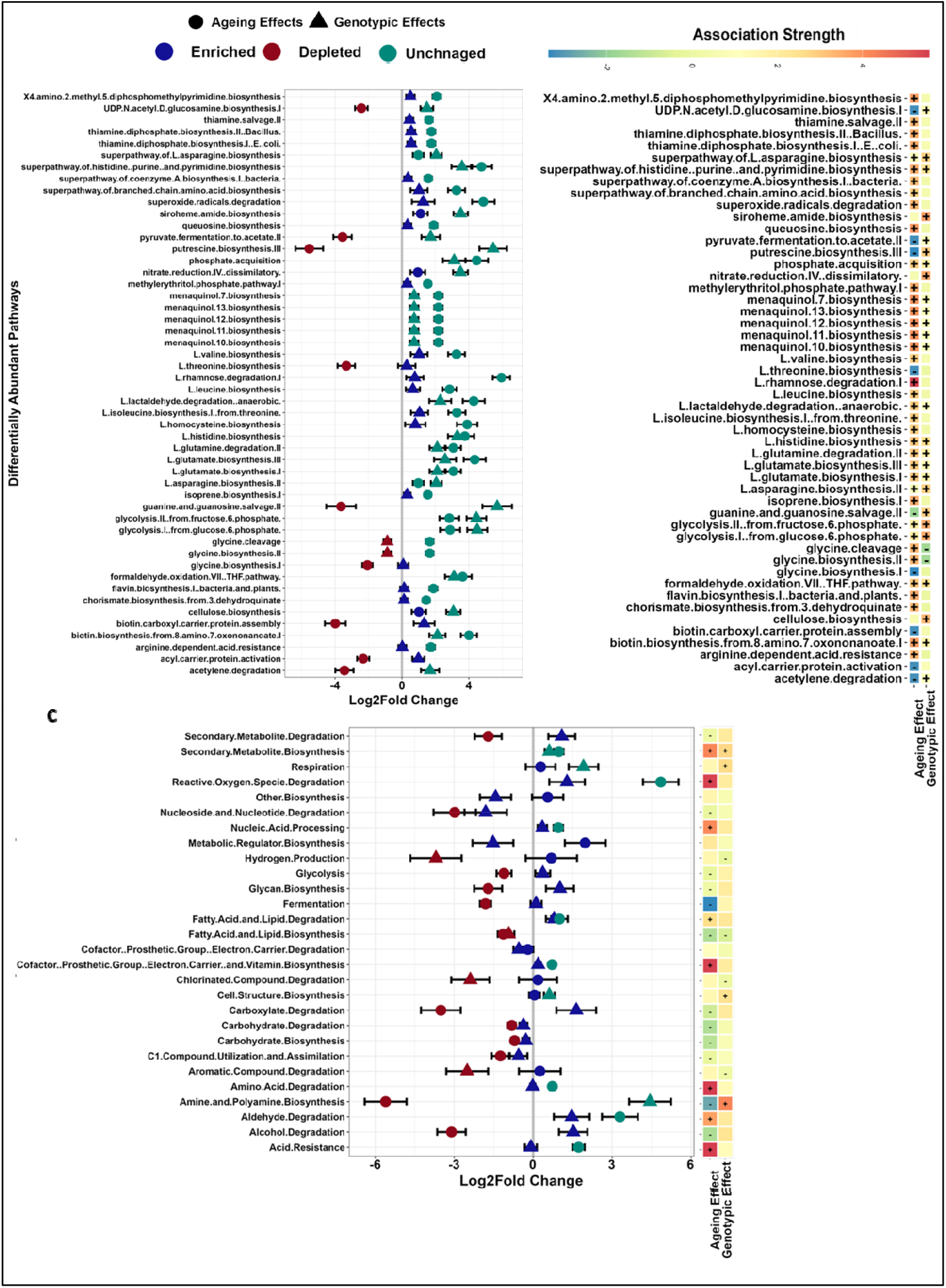
Synuclein pathology triggers distinct functional changes in the gut microbiota. A) Differential abundance plot highlighting predicted microbial pathways being altered due to ageing and genotype in hαSyn-Tg mice. The circles and the squares represent the predicted microbial pathways alterations associated with ageing and genotypic effects respectively. Blue and red colour represents predicted microbial pathways significantly uperegulated (p-value <0.05) and significantly downregulated (p-value <= 0.05) in hαSyn-Tg mice as compared to the WT. Green color represents predicted microbial pathways that are statistically non-significant (p value>0.05)in the hαSyn-Tg mice as compared to the WT. B) Heat-map of the correlation between the predicted microbial pathways abundance in hαSyn-Tg mice with ageing and genotypic effects. Positive and negative significant predictive microbial pathway correlations (p-values <0.05) have been indicated with + and – signs respectively. C) Differential abundance of microbial pathways grouped based on combined with heat map of the the correlation between the predicted microbial pathways abundance in hαSyn-Tg mice with ageing and genotypic effects. Positive and negative significant predictive microbial pathway correlations (p-values <0.05) have been indicated with + and – signs respectively.

### Synuclein pathology triggers distinct functional changes in the gut microbiota

Similar to microbial composition, predicted pathways are also associated with either synuclein accumulation or ageing. Functional pathways associated with fatty acid and lipid biosynthesis are negatively associated with both ageing and genotype, signifying their downregulation with increasing age and presence of transgene. On contrary, a reverse trend is observed in the fatty acid and lipid degradation pathways, reflecting catabolic pathways are favoured enrich with age and presence of the transgene. Pathways associated with secondary metabolite biosynthesis are positively correlated with both ageing and genotypic effects. These pathways are enriched with age and the presence of the transgene. The pathways associated with aldehyde degradation also enriches with both ageing and genotypic effects. Pathways associated with the reactive oxygen species degradation enrich with both age and presence of the transgene but show a weak positive correlation with both ageing and genotypic effects. Pathways involved in glycan production shows a negative correlation with age and a weak positive correlation with genotypic effects as a result of which the pathway is enriched with the presence of the transgene but downregulated with age. A reverse trend is observed with the amine and polyamine biosynthesis pathway; hence, amine and polyamine biosynthesis decrease with age but increase with accumulation of α synuclein. A similar trend is followed by the pathways that are associated with alcohol degradation.

## Discussion

Parkinson’s disease is a progressive neurodegenerative characterized by the accumulation of α-synuclein in the substantia nigra and the loss of dopaminergic neurons. Despite decades of research, the precise aetiology of PD remains elusive. In the recent years, the gut microbiome has has emerged as a critical factor influencing the pathogenesis of the disease. Our findings contribute to the growing body of evidence highlighting the role of the gut-brain axis in PD, specifically examining the relationship between gut dysbiosis and α-synuclein pathology in an animal model of the disease. This study aimed to determine how gut microbial alterations in human A53T α-synuclein-overexpressing transgenic (hαSyn-Tg) mice recapitulate microbiome disturbances observed in PD patients.

We observed gut microbial dysbiosis in hαSyn-Tg mice before the onset of motor symptoms, suggesting that early alterations in microbiota may contribute to prodromal changes associated with PD. Notably, early dysbiosis was marked by an increased abundance of *Staphylococcus* species. Members of the *Staphylococcus* genera are known to produce functional amyloid proteins called phenol soluble modulins, which can promote bacterial virulence and catalyze α-synuclein aggregation [34]. The translocation of these amyloid proteins to the bloodstream and intestinal lumen especially in the presence of increased gut permeability, could exacerbate neuroinflammation and neurodegeneration. Conversely, a decrease in the abundance of *Parasutterella, Lactobacillus and Bifidobacterium* genera was observed before the onset of motor symptoms.

Meta-analysis of clinical PD cohorts have consistently identified depletion of the family Lachnospiraceae as a hallmark of PD-related gut dysbiosis PD. However, in our transgenic mouse model, *Lachnospiraceae* was enriched before motor symptom onset, contrasting with human findings.[35, 36]. Additionally, we observed a reduction in *lactobacillus* and *bifidobacterium species* in the haSyn-Tg mice pre and post-onset of the motor symptoms, suggesting that alpha synuclein aggregation may directly or indirectly inhibit the growth of beneficial bacteria. Similar observations have been recorded in the line61 transgenic mouse model of PD[37]. Interestingly, human PD cohort studies have reported an increased abundance of *Lactobacillus* species, which contrasts with observations in mouse models [38, 39]. *Lactobacillus* and *Bifidobacterium* are commonly used as probiotics and *Lactobacillius* have been shown to reduce dopaminergic neuronal loss in MPTP induced PD mouse model[40].

In healthy individuals, the gut microbiota is typically dominated by the *Firmicutes* and *Bacteroidetes* phyla, which together constitute approximately 90-95% of the colonic microbiome [41]. In our dataset, hαSyn-Tg mice exhibited an increased Firmicutes-to-Bacteroidetes (F/B) ratio before motor symptom onset compared to wild-type (WT) mice. However, as disease progressed, this ratio declined. In contrast, human studies have reported an increasing F/B ratio with aging[42]. A reduction in Bacteroidetes population is often associated with a decrease production of short chain fatty acid (SCFAs particularly butyrate and propionate [39]. SCFAs, play a crucial role in maintaining gut homeostasis, reducing inflammation, and supporting gut barrier integrity. Studies in α-synuclein mouse models consistently report a decline in SCFA-producing bacteria, which may contribute to weakened gut barrier function, increased systemic inflammation, and accelerated α-synuclein aggregation and neuroinflammation. Notably, SCFA supplementation, particularly butyrate, has been shown to mitigate neuroinflammation, reduce α-synuclein pathology, and improve motor symptoms in PD models [14].

Interventions that modulate gut microbiota have demonstrated potential in alleviating PD-related pathology in α-synuclein mouse models. The administration of broad-spectrum antibiotics to alter gut microbiota composition has been shown to reduce motor deficits and α-synuclein accumulation in the brain, highlighting a direct link between gut dysbiosis and PD-like pathology [43]. Likewise, probiotic supplementation aimed at restoring a healthy gut flora has reported to reduce neuroinflammation and improve motor symptoms, likely by modulating the gut microbiota composition and enhancing SCFA production.

Our study also identified a higher abundance of *Akkermansia muciniphila* in aged transgenic mice, reflecting dynamic microbial population shifts. *A. muciniphila* is a mucus-degrading bacterium associated with increased gut permeability, and its elevated abundance in PD models may indicate greater susceptibility to epithelial barrier dysfunction, ultimately compromising gut integrity [44].

Our study identifies significant gut microbial changes in hαSyn-Tg mice both before and after the onset of motor symptoms. We observed alterations in gut microbiota composition associated with both aging and α-synuclein aggregation, highlighting potential links between gut dysbiosis and PD pathology. While this model recapitulates some aspects of gut microbial alterations observed in human PD patients, it does not fully capture all human-specific microbiome changes. These discrepancies may be due to inherent differences in gut microbial composition between humans and mice [47]. Furthermore, most associations identified in human and mouse PD models remain correlative rather than causal.

Future studies employing faecal microbiota transplantation (FMT) from PD patients with gut dysfunction into germ-free mice could help establish a causal relationship between gut dysbiosis and PD progression. Such experiments would provide deeper mechanistic insights into the functional role of the gut microbiome in PD pathogenesis. Understanding these microbial interactions may open new avenues for microbiome-targeted therapeutic interventions aimed at delaying or preventing PD progression.

**Table 1.1:**
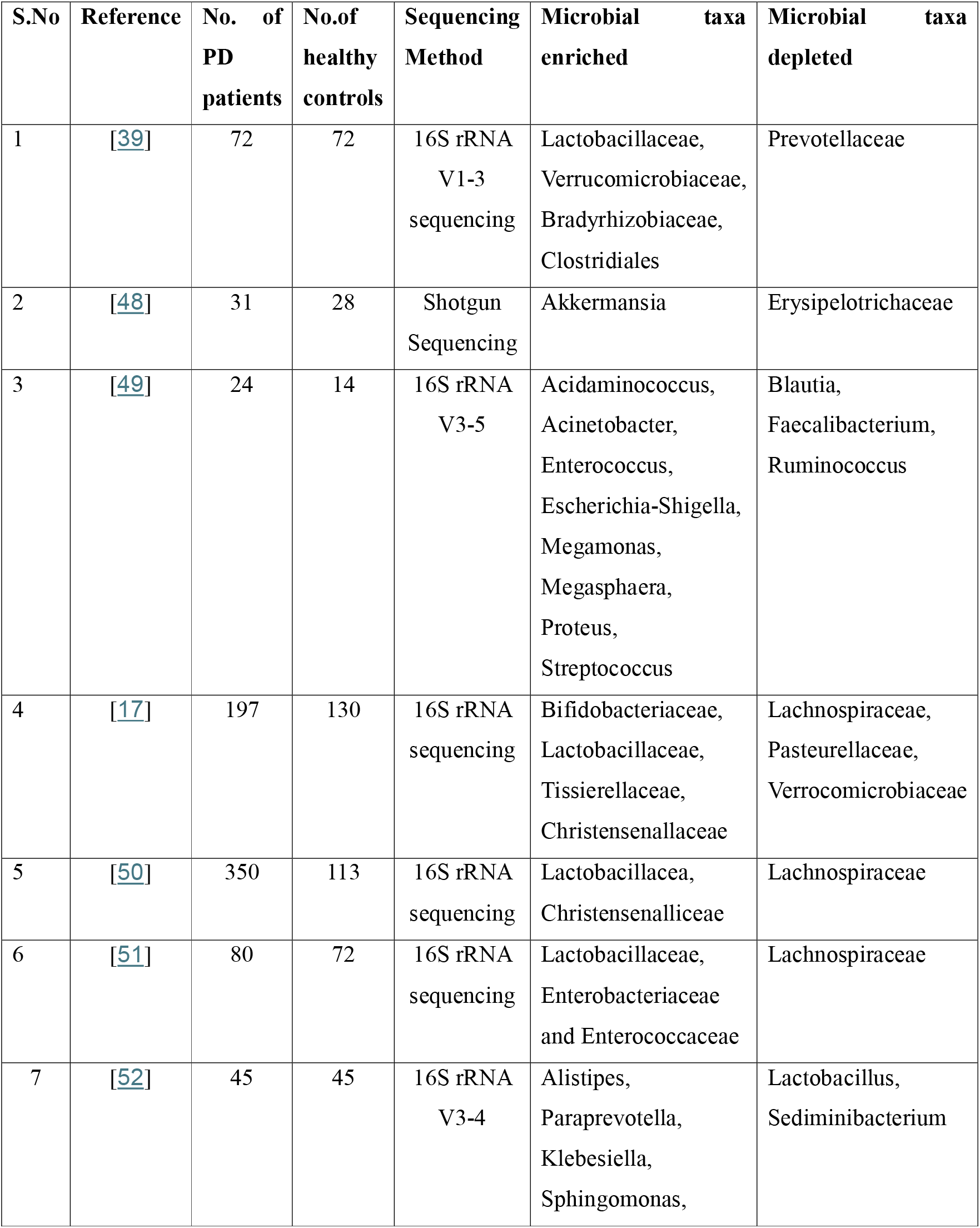
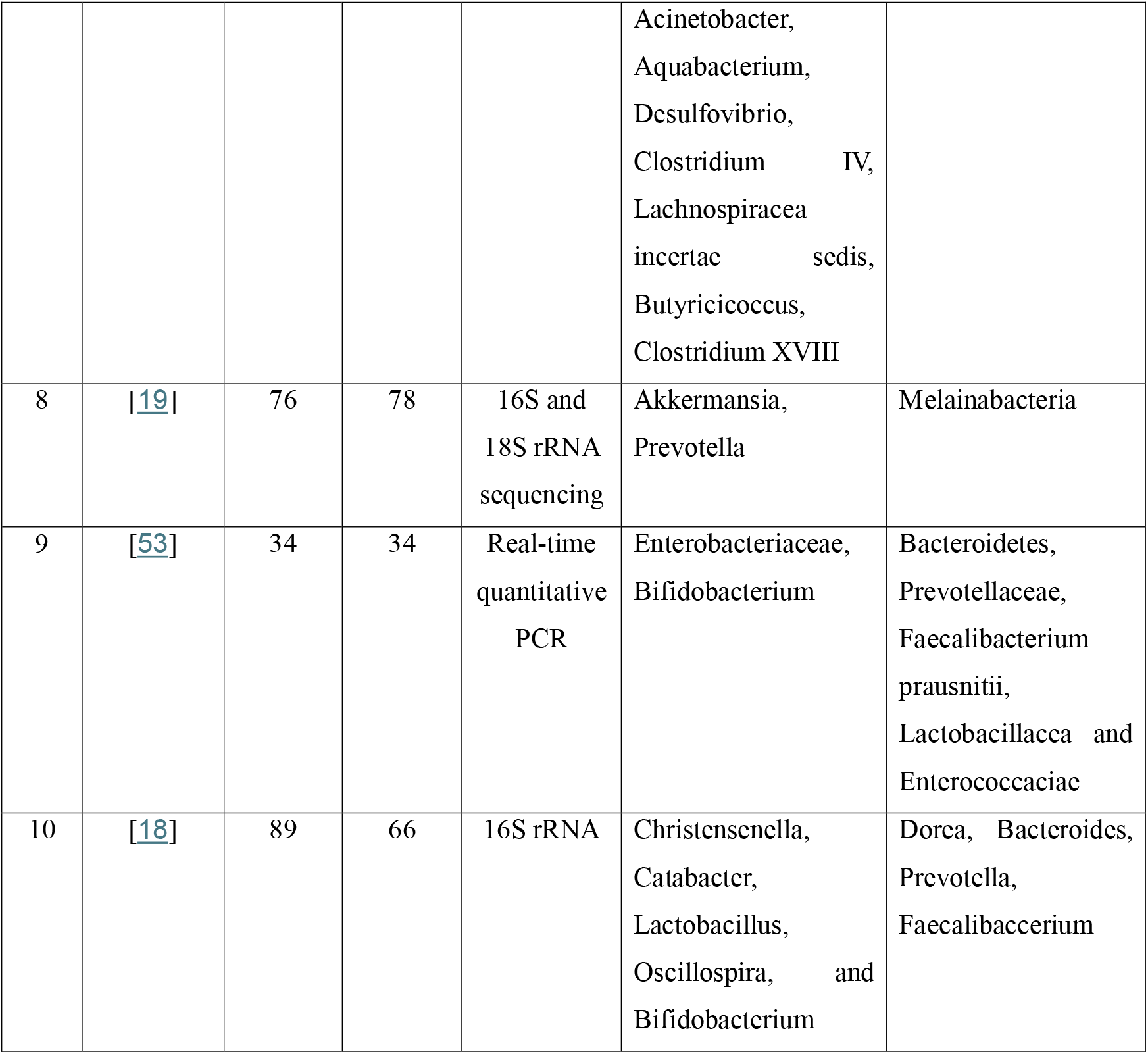
Gut microbial alterations identified across different Parkinson’s cohorts.

**Table 1.2:** Gut microbial alterations identified across different animal models of Parkinson’s disease.

| S.No | Reference | Animal Model | Sample Size | Sequencing Method | Microbial Taxa enriched | Microbial Taxa depleted |
| --- | --- | --- | --- | --- | --- | --- |
| 1 | [54] | Rotenone induced C57BL/6 Mice | N=5 | 16SrRNA V4-5 sequencing | Lactobacillus | Bacteroidetes, Clostridium and Sutterella |
| 2 | [55] | Mitopark<br>Transgenic<br>Mice | N=8<br>(PD) and<br>N=10<br>(non<br>PD) | 16SrRNA<br>V3-4<br>sequencing | Prevotella |  |
| 3 | [37] | Thy1-hSYN<br>“Line 61”<br>Mice | N=28<br>(TG) and<br>32<br>(NTG) | Shotgun<br>Sequencing |  | Lactobacillus<br>and<br>Bifidobacterium |

## Supporting information

Metadata

## Acknowledgements

This research has been made possible through the support of the US Department of Defence, the Michael J. Fox Foundation for Parkinson’s Disease National health and medical research council (NHMRC) of Australia, Shake It Up Australia, and the funding from University of Queensland awarded to Dr. Richard Gordon. We also wish to thank Queensland University of Technology and Translational research institute for all the resources that facilitated this work.

## Data availability

All datasets generated and analysed are included in the supplementary files.

## Code availability

All codes used for analysis are included in the supplementary files.

